# ARBitR: An overlap-aware genome assembly scaffolder for linked reads

**DOI:** 10.1101/2020.04.29.065847

**Authors:** Markus Hiltunen, Martin Ryberg, Hanna Johannesson

## Abstract

10X Genomics Chromium linked reads contain information that can be used to link sequences together into scaffolds in draft genome assemblies. Existing software for this purpose perform the scaffolding by joining sequences together with a gap between them, not considering potential contig overlaps. Such overlaps can be particularly prominent in genome drafts assembled from long-read sequencing data where an overlap-layout-consensus (OLC) algorithm has been used. Ignoring overlapping contig ends may result in genes and other features being incomplete or fragmented in the resulting scaffolds. We developed the application ARBitR to generate scaffolds from genome drafts using 10X Chromium data, with a focus on minimizing the number of gaps in resulting scaffolds by incorporating an OLC step to resolve junctions between linked contigs. We tested the performance of ARBitR on three published and simulated datasets and compared to the previously published tools ARCS and ARKS. The results revealed that ARBitR performed similarly considering contiguity statistics, and the advantage of the overlapping step was revealed by fewer long and short variants in ARBitR produced scaffolds, in addition to a higher proportion of completely assembled LTR retrotransposons. We expect ARBitR to have broad applicability in genome assembly projects that utilize 10X Chromium linked reads.

**Availability and implementation:** ARBitR is written and implemented in Python3 for Unix-like operative systems. All source code is available at https://github.com/markhilt/ARBitR under the GNU General Public License v3.

**Supplementary information:** *available online*

## 1 Introduction

Contiguity in genome assemblies is important for the ability to analyze e.g. structural rearrangements, gene order, synteny between divergent genomes, linkage between genetic variants, and repetitive genomic regions. A few years ago, 10X Genomics launched the Chromium system, which leverages microfluidics to partition DNA molecules into droplets where unique barcode sequences are attached. Libraries are subsequently sequenced using Illumina short-read sequencing. This way, reads originating from adjacent regions in the genome will share a high proportion of barcodes, providing linkage information of the reads for ranges up to the length of the input DNA molecule. Such information can be used to order previously assembled contigs into scaffolds (Sedlazeck *et al*., 2018; Yeo *et al*., 2017).

There are published pipelines designed for genomic scaffolding using 10X Chromium linked reads. To infer linkage of draft contigs, either ARCS or ARKS can be used (Yeo *et al*., 2017; Coombe *et al*., 2018). ARCS finds linkage based on barcoded read mappings while ARKS instead uses a kmer-based approach for finding reads that share a barcode among unmapped read sets, reducing processing time by skipping the mapping step (Yeo *et al*., 2017; Coombe *et al*., 2018). After linkage of contigs has been determined by ARCS or ARKS, LINKS is used to merge the linked sequences by inserting a gap between them (Warren *et al*., 2015). While ARCS/ARKS are able to estimate a gap distance between putative linked contigs, LINKS does not resolve cases where there is an estimated negative distance, i.e. an overlap between contigs, and simply merges them with a single lower-case “n” in between. Including a gap in the merged sequence leads to the risk of fragmenting genes and other features in those regions. Especially for genomes where repeat clusters are relatively short and the draft assembly was produced by an overlap-layout-consensus (OLC) algorithm (Miller *et al*., 2010), such overlaps may be quite frequent. Instead of indiscriminately merging the contigs with a gap, the scaffolding could be performed by using OLC. In an attempt to minimize the number of gaps while building scaffolds based on 10X Chromium linked-read information, we developed the tool ARBitR: Assembly Improvement using Linked reads. Designed for flexibility while requiring minimal user input, ARBitR performs the linkage-finding and scaffolding steps in succession in a single Python application.

## 2 Materials and Methods

ARBitR relies on 10X Chromium barcode information to determine linkage of input contigs (Figure 1). To collect barcodes, ARBitR utilizes the Pysam module (https://github.com/pysam-developers/pysam) which acts as a wrapper for samtools (Li *et al*., 2009). Links are computed and represented in a graphical data structure that we refer to as a linkgraph. Paths through the graph are determined, and at each step in the path, ARBitR detects overlaps between the involved sequences using the Python module Mappy (Li, 2018, 2). If no overlap is found, ARBitR inserts a gap between the sequences similarly to LINKS.

**Figure 1:**
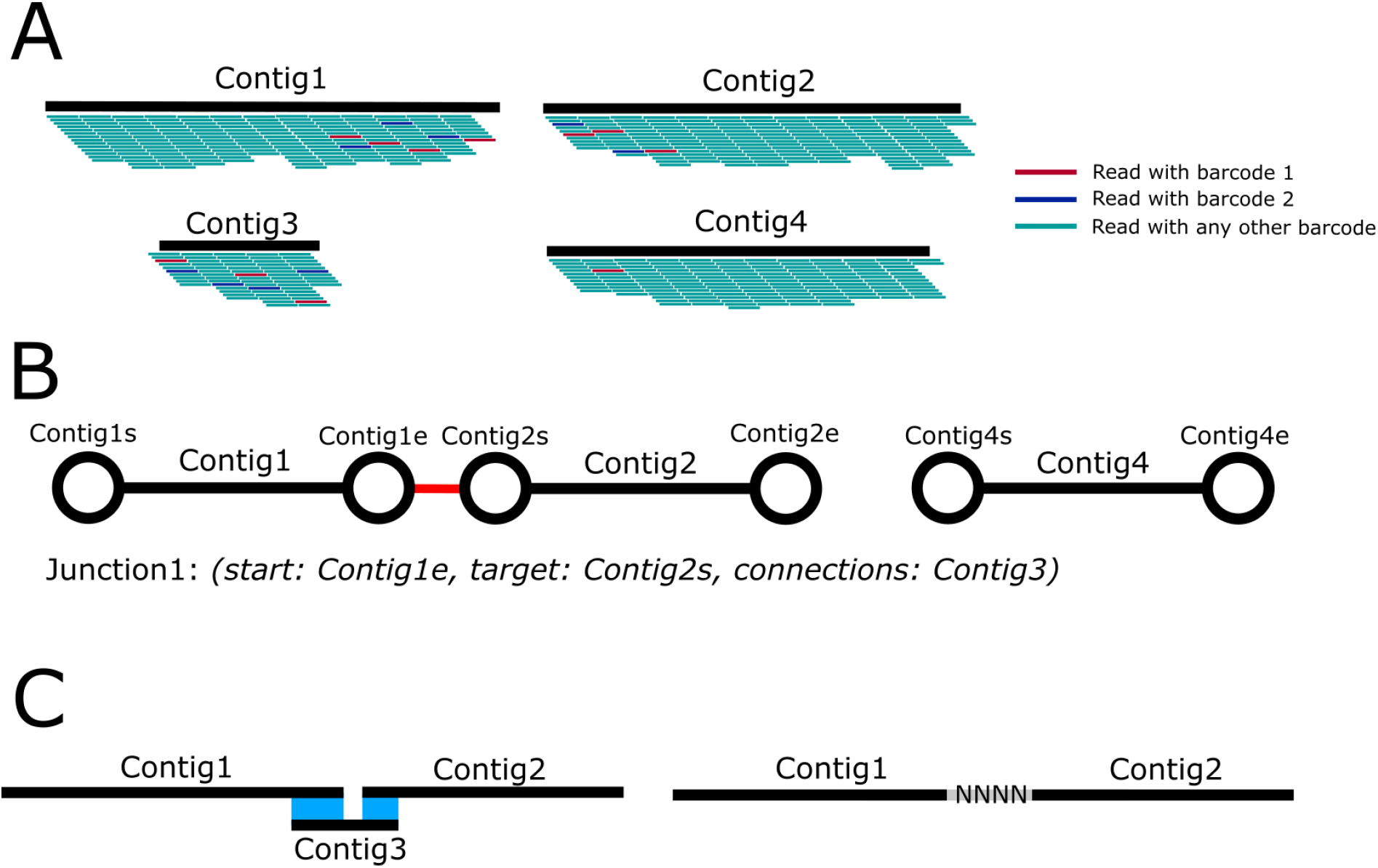
ARBitR pipeline. A. To infer linkage of contigs in the input genome assembly, ARBitR relies on GEM barcode information of 10X Chromium linked reads that have been mapped to a draft genome assembly. Short input contigs (contig3) often consist of repetitive sequence and may obstruct efficient scaffolding because of unreliable read mapping, and are initially disregarded. From the starting (s) and ending (e) regions of the long contigs, GEM barcodes are collected. For each region, the fraction of shared barcodes with every other region is computed, and significant outliers are determined. B. The outliers are collected and represented in a graphical format. Nodes in the graph correspond to input sequence start and end regions, and edges correspond to outlying fractions of shared barcodes between these regions. The barcodes at each edge are kept in memory. The graph is output in the Graphical Fragment Assembly format (gfa), viewable with e.g. Bandage (Holt *et al*., 2015). The paths through the graph that can be unambiguously traversed are then determined. At each step in the path, termed junction, ARBitR searches the short input contigs for ones that share a high fraction of barcodes with the junction. Such short contigs potentially originated from the genomic region in question. C. Finally, sequences are produced from the paths, by first trimming input sequences for regions of low read coverage, after which every junction in every path is visited and resolved by overlap-layout-consensus (OLC). If no complete path through the junction can be found by OLC, as many sequences as possible are included and a gap is inserted where no overlap can be found between sequences.

To test the performance of ARBitR in relation to the ARCS/ARKS and LINKS pipeline, we utilized three datasets: (1) published PacBio, Nanopore and 10X Chromium linked reads of the fungus *Marasmius oreades* (Hiltunen *et al*., 2019), (2) publicly available PacBio and 10X Chromium reads of *Arabidopsis thaliana* (Sun *et al*., 2019; Jiao and Schneeberger, 2019) and (3) simulated PacBio and 10X Chromium data of *Caenorhabditis elegans*. For dataset 3, we used PBSIM and LRSIM to simulate PacBio and 10X Chromium linked reads, respectively, from the reference genome PRJNA275000 (Asai *et al*., 2012; Luo *et al*., 2017; Thompson *et al*., 2015).

For each of the three datasets, long reads were assembled with Canu (Koren *et al*., 2017), and linked read sets were mapped to each respective draft assembly using either BWA mem or EMA (Li and Durbin, 2009; Shajii *et al*., 2018). ARBitR was used to create scaffolds from the three assemblies. For comparison, we also used ARCS v1.1.1 both in default and in ARKS mode in combination with LINKS v1.8.6 to create scaffolds. Assembly statistics were collected with Quast v5.0.2 (Gurevich *et al*., 2013), using reference genomes for comparison (Maror1, TAIR10 for datasets 1 and 2, respectively) (Berardini *et al*., 2015; Hiltunen *et al*., 2019). Reasoning that higher-quality assemblies should have fewer variants compared to the 10X Chromium reads than assemblies of lower quality, we utilized the Longranger WGS pipeline for variant calling (https://www.10xgenomics.com). As contigs often break during assembly in regions where long terminal repeat (LTR) retrotransposons are found, we additionally estimated the number of complete and fragmented LTR elements using the LTR Assembly Index statistic (LAI) (Ellinghaus *et al*., 2008; Ou *et al*., 2018; Ou and Jiang, 2018). Datasets and software parameters are described in detail in Supplementary Methods and Supplementary Table S1. Computations were performed on a Dell server on Ubuntu 18.04.3 using a maximum of 48 cores and with 503 Gb available memory.

## 3 Results

Detailed statistics of the performance of the different scaffolding procedures can be viewed in Supplementary Table S2. In brief, ARBitR found a higher number of linked contigs in the *M. oreades* and *C. elegans* datasets than the other pipelines, resulting in assemblies with fewer scaffolds, higher N50 and lower L50 values, in addition to the longest scaffolds most closely resembling the reference scaffold lengths. In *A. thaliana*, ARKS produced the most contiguous assembly. ARBitR scaffolds from all datasets contain the fewest base mismatches and indels when comparing to reference assemblies, with the only exception of the *C. elegans* raw assembly which has fewer mismatches. The Longranger variant calling pipeline revealed the fewest variants in ARBitR-scaffolded genomes in most cases. Finally, the LAI is higher in ARBitR scaffolds, reflecting the advantage of overlap-based scaffolding for assembling LTR elements. In the case of *M. oreades*, the reference assembly is fragmented, and in the ARBitR scaffolds of this genome we find a higher number of assembled LTRs than in the reference. We noticed instances where genomic features appear fragmented or duplicated in LINKS scaffolds while being more complete in overlap merges performed by ARBitR (Figure S1).

## 4 Conclusion

We present a new method, ARBitR, to apply 10X Chromium linked-read information for scaffolding of draft genome assemblies. In comparing contiguity statistics of our tested datasets, ARBitR performs better than, or similarly to, previously published pipelines. A key feature of the ARBitR pipeline is the consideration of overlaps between ends of linked contigs, which we found can decrease the number of indels and mismatches in resulting scaffolds and improve assembly of transposable elements. We expect ARBitR to have broad applicability in genome assembly projects that utilize 10X Chromium linked reads, particularly in cases where repeat clusters are relatively short and the initial assembly was produced by an OLC-based algorithm.

