## Supplementary information for "ARBitR: An overlap-aware genome assembly scaffolder for linked reads"

### Method details

*Assembly pre-processing*

As a high-confidence reference genome of *Marasmius oreades* that could be used to quality control our raw genome assembly has yet to be published, we subjected our raw assembly to an initial quality checkup using Tigmint prior to scaffolding (Jackman *et al.*, 2018). Tigmint uses 10X Chromium linked reads to spot regions in draft assemblies where the barcode coverage drops below a certain threshold, as such regions potentially have been wrongly assembled. We used tigmint-molecule to spot such regions (-s 10000) and custom scripts to break the assembly where such regions were found (available at https://github.com/markhilt/genome_analysis_tools).

*Longranger variant calling*

As an additional measurement of genome concordance, we characterized variation between each assembly and the 10X Chromium reads, with the assumption that better assemblies have fewer called variants. For this purpose, we applied the Longranger WGS pipeline to all assemblies, for *M. oreades* and *A. thaliana* we used Longranger v2.2.2 and for *C. elegans* v2.1.4, as the newer version was incompatible with simulated reads. We quantified the number of calls with a PASS quality flag for large-scale structural variations, mid-scale deletions and single-nucleotide variants in addition to short indels and summarized in Table S2.

*Long Terminal Repeat analysis*

As genome assemblies are known to break in repetitive regions, and Long Terminal Repeat (LTR) transposable elements are the most common repeats in plant and fungal genomes (Ou and Jiang, 2018; Castanera *et al.*, 2016), we quantified the extent to which LTRs were assembled in the *M. oreades* and *A. thaliana* genomes (the low amount of LTR elements in the *C. elegans* genome made this analysis impossible in this case). The analysis was performed as recommended by the authors of LTR_retriever (Ou and Jiang, 2018): first find candidates of LTR elements in the genome using LTRharvest (-minlenltr 100 -maxlenltr 7000 -mintsd 4 -maxtsd 6 -motif TGCA -motifmis 1 -similar 85 -vic 10 -seed 20 -seqids yes) and LTR_finder (Ellinghaus *et al.*, 2008; Xu and Wang, 2007), then use LTR_retriever to quantify how common these elements are in the genome and calculate the LTR Assembly Index.

### Supplementary tables

**Table S1**. Datasets used for benchmarking ARBitR, ARCS and ARKS.

| Species | Long read data source | 10X Chromium data source | Read mapper | ARBitR parameters | ARCS parameters | ARKS parameters | Data reference |
|---|---|---|---|---|---|---|---|
| *Marasmius oreades* | SRA: PRJNA525964 | SRA: PRJNA525964 | EMA | -s25000 -Q15 -f10 -F0 -b5 -c5 | -e25000 | default | (Hiltunen *et al.*, 2019) |
| *Arabidopsis thaliana* | SRA: ERR3415826 | ENA: ERR2851508 | BWA mem | -s35000 -m50000 -Q15 -F0.03 -f39 | default | -D | (Sun *et al.*, 2019) |
| *Caenorhabditis elegans* | Simulated (PBSIM) | Simulated (LRSIM) | EMA | -s45000 -m50000 -Q20 -b4 -c0 | -c8 | -D | (Thompson *et al.*, 2015) |

**Table S2.** Scaffolding performance of ARBitR, ARCS and ARKS. Bold font indicates values closest to the references for each parameter.

| Species | Assembly | # Scaffolds | N50 (bp) | L50 | Longest scaffold (bp) | Mismatches /100kb | Indels/100kb | Longranger SVs | Longranger mid-scale deletions | Longranger SNVs & short indels | LAI[a] |
|---|---|---|---|---|---|---|---|---|---|---|---|
| *M. oreades* | Maror1[b] | 39 | 3564693 | 6 | 5044703 | NA | NA | 0 | 31 | 6412 | 8.82 |
| | Raw assembly | 163 | 733082 | 18 | 2593343 | 17.17 | 71.77 | **0** | 111 | 29011 | 7.59 |
| | ARBitR | **69** | **2560603** | **7** | **4675370** | **12.66** | **63.08** | **0** | **107** | **28850** | 10.37 |
| | ARCS | 108 | 1501983 | 9 | 4619992 | 17.50 | 71.84 | **0** | 123 | 28915 | 7.51 |
| | ARKS | 122 | 1163884 | 12 | 2953429 | 17.20 | 71.83 | **0** | 108 | 28961 | **7.61** |
| *A. thaliana* | TAIR10[b] | 7 | 23459830 | 3 | 30427671 | NA | NA | 2 | 849 | 877989 | 17.40 |
| | Raw assembly | 771 | 454218 | 79 | 2359306 | 905.48 | 298.23 | 28 | 1243 | 1089317 | 7.59 |
| | ARBitR | 472 | 4635812 | 10 | 10687703 | **900.75** | **295.56** | **2** | 1214 | **1082913** | **17.36** |
| | ARCS | **433** | 4354541 | 9 | 12501495 | 902.43 | 298.42 | 4 | 1183 | 1085541 | 6.63 |
| | ARKS | 441 | **6950158** | **7** | **12820601** | 904.76 | 298.32 | 13 | **1180** | 1085071 | 6.87 |
| *C. elegans* | PRJNA275000[b] | 7 | 17183857 | 3 | 20182852 | NA | NA | 6 | 0 | 6494 | NA |
| | Raw assembly | 66 | 5629213 | 7 | 12209780 | **1.74** | 3.30 | 82 | 2 | 8479 | NA |
| | ARBitR | **50** | **10218938** | **4** | **20110797** | 1.88 | **3.21** | **42** | **0** | 8483 | NA |
| | ARCS | 54 | 8544338 | 5 | 13042304 | 1.97 | 3.29 | 60 | 2 | **8423** | NA |
| | ARKS | 53 | 9420178 | **4** | 21146662 | 1.74 | 3.31 | 84 | 2 | 8458 | NA |

[a]Long terminal repeat element Assembly Index

[b]Reference genome

### Supplementary figures

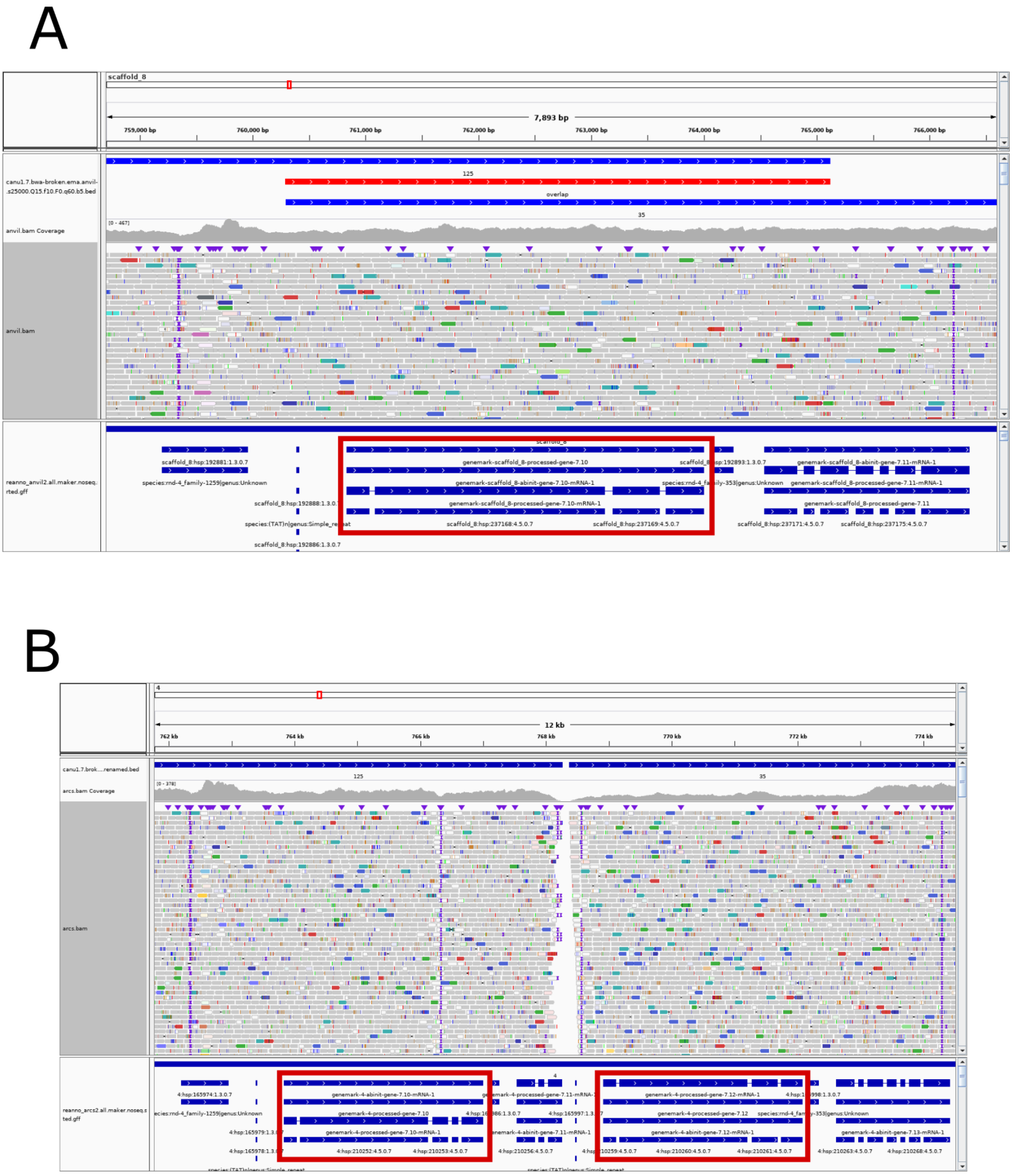


**Figure S1: Examples of scaffolding results.** IGV plots of (A) ARBitR and (B) ARCS & LINKS scaffolding results in a region where ARBitR performed an overlap merge in the *Marasmius oreades* dataset. Tracks from top to bottom: original scaffolds (overlap in red), mapped read coverage, read alignments, gene annotation. Highlighted: duplicated gene in the ARCS assembly. Note also the reduced read coverage in these regions in the ARCS assembly.
